# Hippo signaling promotes Ets21c-dependent apical cell extrusion in the *Drosophila* wing disc

**DOI:** 10.1101/2020.03.03.975771

**Authors:** Xianlong Ai, Dan Wang, Junzheng Zhang, Jie Shen

## Abstract

Cell extrusion is a crucial regulator of epithelial tissue development and homeostasis. Epithelial cells undergoing apoptosis, bearing pathological mutations, and possessing developmental defects are actively extruded toward elimination. However, the molecular mechanisms of *Drosophila* epithelial cell extrusion are not fully understood. Here, we report that activation of the conserved Hippo (Hpo) signaling pathway induces both apical and basal cell extrusion in the *Drosophila* wing disc epithelia. We show that canonical Yorki targets Diap1, and that dMyc and Cyclin E are not required for either apical or basal cell extrusion induced by activation of this pathway. Another target gene, *bantam*, is only involved in basal cell extrusion, suggesting novel Hpo-regulated apical cell extrusion mechanisms. Using RNA-Seq analysis, we found that JNK signaling is activated in the extruding cells. We provide genetic evidence that JNK signaling activation is both sufficient and necessary for Hpo-regulated cell extrusion. Furthermore, we demonstrate that the ETS-domain transcription factor Ets21c, an ortholog of proto-oncogenes *FLI1* and *ERG*, acts downstream of JNK signaling to mediate apical cell extrusion. Our findings reveal a novel molecular link between Hpo signaling and cell extrusion.

**SUMMARY STATEMENT:** Activation of Hippo signaling induces cell extrusion in the *Drosophila* wing epithelia, in which *bantam* mediates basal cell extrusion and Ets21c mediates apical cell extrusion.

## INTRODUCTION

The Hippo (Hpo) pathway is conserved from *Drosophila* to mammals (Dong et al., 2007; Pan, 2010). Upstream key components of this pathway, including Hpo and Warts (Wts), form a core kinase cascade to regulate the transcriptional coactivator Yorkie (Yki) (Huang et al., 2005; Praskova et al., 2008; Udan et al., 2003; Wu et al., 2003). When the Hpo pathway is inactive, dephosphorylated Yki is translocated into the nucleus and interacts with the DNA-binding transcription factor Scalloped (Sd) (Zhang et al., 2008). The Yki-Sd complex induces the expression of a wide range of target genes, including the *Cyclin E* (*CycE*) and *dMyc* cell-cycle regulators, *bantam* (*ban*) microRNA, and *Drosophila inhibitor of apoptosis protein 1* (*Diap1*) (Pan, 2010). Numerous studies have indicated that the Hpo pathway functions as a tumor suppressor. Yap is hyperactivated in some human cancers, including lung cancer (Yang et al., 2010) and uveal melanoma (Feng et al., 2014; Yu et al., 2014). However, YAP is also downregulated in individuals with multiple myeloma and in those with acute myeloid leukemia (Cottini et al., 2014).

The Hpo pathway is essential for epithelial tissue development and homeostasis by regulating cell survival (Tapon et al., 2002), cell proliferation (Halder and Johnson, 2011), organ size (such as that of the *Drosophila* eye and thorax and the mouse liver) (Dong et al., 2007), and cell-cell adhesion (Schroeder and Halder, 2012). It has been reported that *yki* mutant clones grow poorly in the third instar larval wing disc (Qing et al., 2014). In addition to roles in apoptosis and cell proliferation, the Hpo pathway has been recently implicated in cell extrusion and invasion, which may help to explain the poor survival of *yki* mutant clones. Expression of *yki*RNAi along the anterior/posterior (A/P) boundary of *Drosophila* wing discs induces cell migration across the A/P boundary and basal cell extrusion (BCE) (Ma et al., 2017). However, except for apoptosis induced BCE, whether cell extrusion causes the abnormal Hpo-Yki signaling-induced poor recovery rate is unclear.

In both invertebrate and vertebrate models, cell extrusion (Gudipaty and Rosenblatt, 2017) plays an important role in epithelial homeostasis and development as well as in cancer cell metastasis (Eisenhoffer et al., 2012; Gu and Rosenblatt, 2012; Gudipaty and Rosenblatt, 2017; Marinari et al., 2012; Ohsawa et al., 2018). Moreover, extruded tumor cells have invasive abilities (Dunn et al., 2018; Pagliarini and Xu, 2003; Shen et al., 2014; Vaughen and Igaki, 2016). Here, we tested the hypothesis that the Hpo pathway maintains tissue homeostasis by suppressing cell extrusion. We found that Hpo pathway activation, by repressing *yki* or expressing *hpo/wts*, induced both apical cell extrusion (ACE) and BCE. Furthermore, *ban* expression could suppress BCE but not ACE. We further demonstrated that JNK signaling was necessary and sufficient to induce ACE downstream of Yki. Mechanistically, we present genetic evidence that Hpo-JNK pathway induced ACE is mediated by Ets21c, a member of the ETS-domain transcription factor family.

## RESULTS

### Activated Hpo pathway induced cell extrusion

This study was designed to investigate whether poor *yki* mutant clone growth is due to the effect of cell extrusion, in addition to cell death and proliferation constriction. We used mosaic analysis with a repressible cell marker (MARCM) (Lee and Luo, 2001) to generate GFP marked *yki* mutant clones and monitored their cellular behavior. Compared with the control (Fig. S1A), wing pouch *yki* mutant clones were rare and small (Fig. S1B). Next, we generated *UAS*-*ykiRNAi* and *UAS*-*GFP* co-expressing clones using flp-out technology (Harrison and Perrimon, 1993). *ykiRNAi* clonal cells vanished when grown for 72 h (Fig. S1C, D). Given that Yki is a cell death suppressor (Huang et al., 2005), the loss of clones may be caused by apoptosis. To address this possibility, *p35*, encoding a pan-caspase inhibitor (Hay et al., 1995), was co-expressed with *ykiRNAi* in the clonal cells to suppress potential apoptosis.

However, even when apoptosis was suppressed by *p35* expression, *ykiRNAi* clones were still very small and rare, especially in the wing pouch (Fig. S1E). Therefore, the low recovery rate of clones lacking Yki activity was not due to apoptosis. These results support the possibility of cell extrusion in *yki* mutant clones.

To check whether cells with downregulated *yki* are extruded from the wing disc, vertical wing disc sections were examined by y-z scanning on a confocal microscope. The apical side of disc proper (DP) cells faces the lumen of the wing sac, and their basal side is the external surface which contacts hemolymph. We used Discs large 1 (Dlg1) (Knust and Bossinger, 2002) as the apical maker of the epithelia during y-z scanning. Compared with wild-type clones (Fig. 1A), *yki* mutant clones rare and invaded into the lumen (Fig.1B, arrow). *ykiRNAi* clones were rare and extruded either basally or apically (Fig. 1C, D). To overcome the low recovery rates during clonal analysis, *UAS*-*ykiRNAi* was expressed under the control of a pan-disc-specific driver *C765*-*Gal4* (Fig. S2A, A’). There were two stripe regions close to the wing hinge folds with greatly reduced Dlg staining in the apical wing disc confocal sections. We inspected the vertical sections of that region and found that there were more cells undergoing ACE when *yki* expression was silenced (Fig. 1E, F, arrow). At the same time, BCE was also evident (Fig. 1E, F, arrowhead). ACE and BCE were observed in over 90% of the wing discs (Fig. S3A, C). We confirmed that the same ACE and BCE phenotypes were observed using another, independent, *UAS*-*ykiRNAi* line (Fig. S2B).

**Fig. 1.**
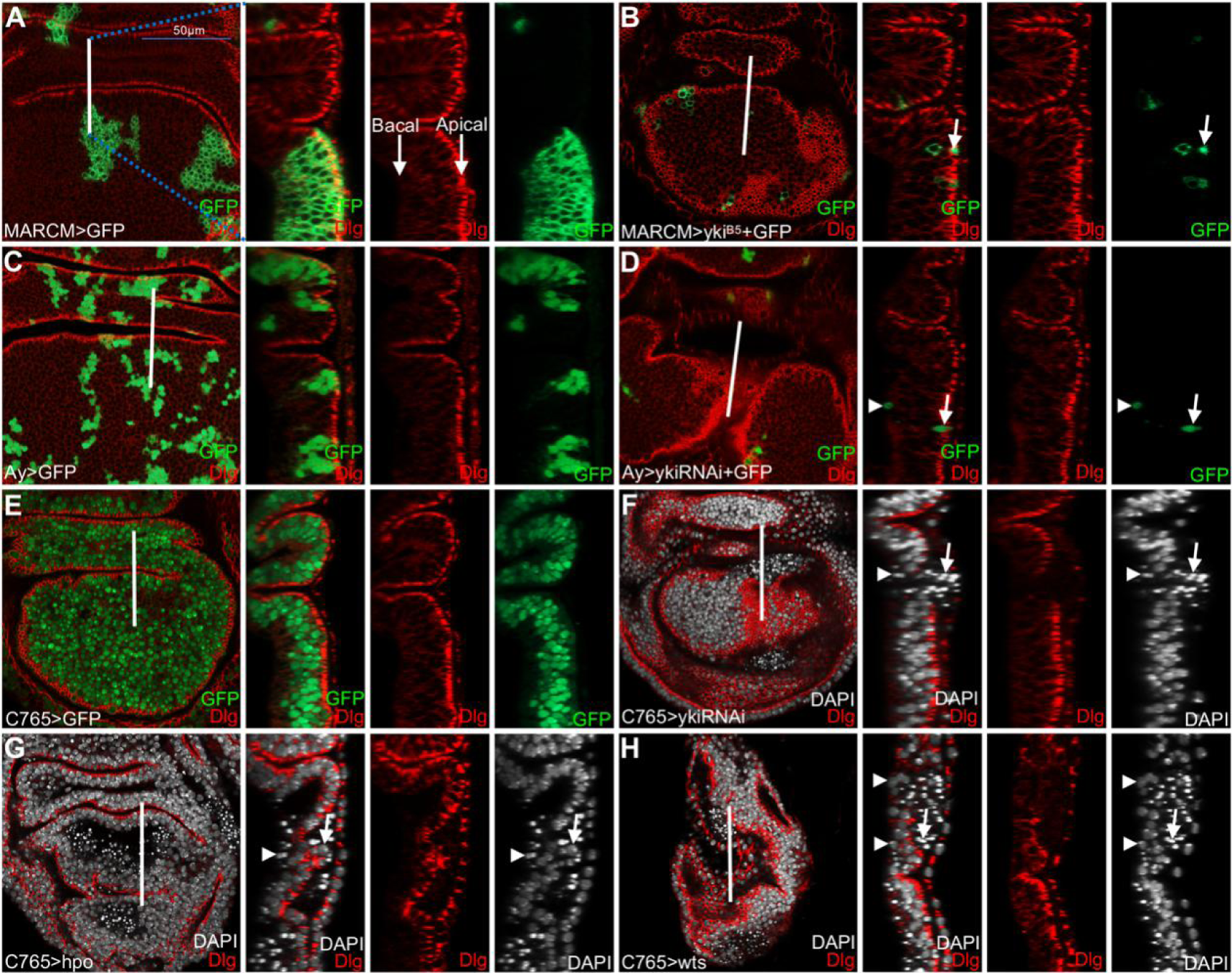
Hpo pathway activation induces ACE and BCE. The developmental stage of the wing imaginal discs shown in this and subsequent figures was middle to late 3^rd^ instar, unless indicated otherwise. To better observe the apically located cells, x-y images of apical sections of the epithelia were scanned by confocal microscopy, unless indicated otherwise. y-z images were scanned at the position indicated by white straight lines and they are oriented with the dorsal side up and the apical side to the right. Dlg staining was used to determine the apical side of the wing epithelia. Arrows indicate apical cell extrusion (ACE) and arrowheads indicate basal cell extrusion (BCE), unless indicated otherwise. The *yki*-depleted line used was *UAS*-*ykiRNAi*^*THU0579*^, unless indicated otherwise. Scale bar is 50 μm. (A) *GFP* control clones generated using the MARCM system. (B) MARCM system-generated *yki*^*B5*^ mutant clones were rare and tended toward apical extrusion (arrow). (C) *GFP* control clones generated using the flp-out system with act5c>*CD2*>Gal4; UAS-GFP. (D) *ykiRNAi* clones were rare and tended to apical extrusion (arrow) or basal extrusion (arrowhead). (E) *GFP* expressing cells driven by *C765*-*Gal4* were not extruded. (F-H) *ykiRNAi* (F), *hpo* (G), and *wts* (H) expressing cells were extruded into the lumen (arrows) and to the basal side of the epithelia (arrowheads).

Hpo and Wts suppress Yki in the *Drosophila* wing disc (Huang et al., 2005). To confirm that activated Hpo signaling regulates cell extrusion, we inspected the vertical sections of wing discs expressing *hpo* and *wts*. Consistently, both ACE and BCE was observed in more than 95% of *UAS*-*hpo*-expressing wing discs (Fig. 1G and Fig. S3). Both ACE and BCE were observed in the *UAS*-*wts*-expressing wing disc pouch at high frequency (76.5% and 100%, for ACE and BCE, respectively, n = 34) (Fig. 1H and Fig. S3). These results suggest that the activated Hpo pathway induces both ACE and BCE in *Drosophila* wing discs. The absence of *yki* expression induces apoptosis (Huang et al., 2005), and this was confirmed by anti-cleaved caspase-3 staining (Fig. S4A). Vertical sections showed that most of the dying cells were located either in the lumen or at the basal side of the epithelia (Fig. S4A). To examine whether ACE and BCE are induced as a side effect of apoptosis, we co-expressed *p35* and *ykiRNAi.* Both ACE and BCE still occurred (Fig. S4B, C) at high frequency in clones co-expressing *p35* and *ykiRNAi* (74.2%, n = 31) (Fig. S3A, C). These data indicate that the Hpo pathway-induced cell extrusion is apoptosis independent.

### The Yki pathway target *ban* mediated BCE

We next tested whether canonical Hpo-Yki target genes mediate the observed cell extrusion. We checked four targets downstream of Yki, including Cyclin E (CycE), a cell cycle regulator (Tapon et al., 2002), Diap1, an apoptosis inhibitor (Wu et al., 2003), *ban*, a microRNA that promotes growth and inhibits cell death (Nolo et al., 2006; Thompson and Cohen, 2006), and dMyc, a critical cellular growth effector (Neto-Silva et al., 2010). Co-expressing *ban* and *ykiRNAi* still induced ACE (Fig. 2A, B) at a high frequency (70%, n = 30) (Fig. S3A and C), but the frequency of BCE in wing discs was reduced (16.7%, n = 30). Co-expressing *Diap1* (Fig. 2C, D), *dMyc* (Fig. 2E, F), and *CycE* (Fig. 2G, H) showed little effect on ACE and BCE frequencies (Fig. S3A and C). These results suggest that silenced-*yki* induced BCE is largely dependent on *ban*, but silenced-*yki* induced ACE is not dependent on the canonical Hpo-Yki targets examined.

**Fig. 2.**
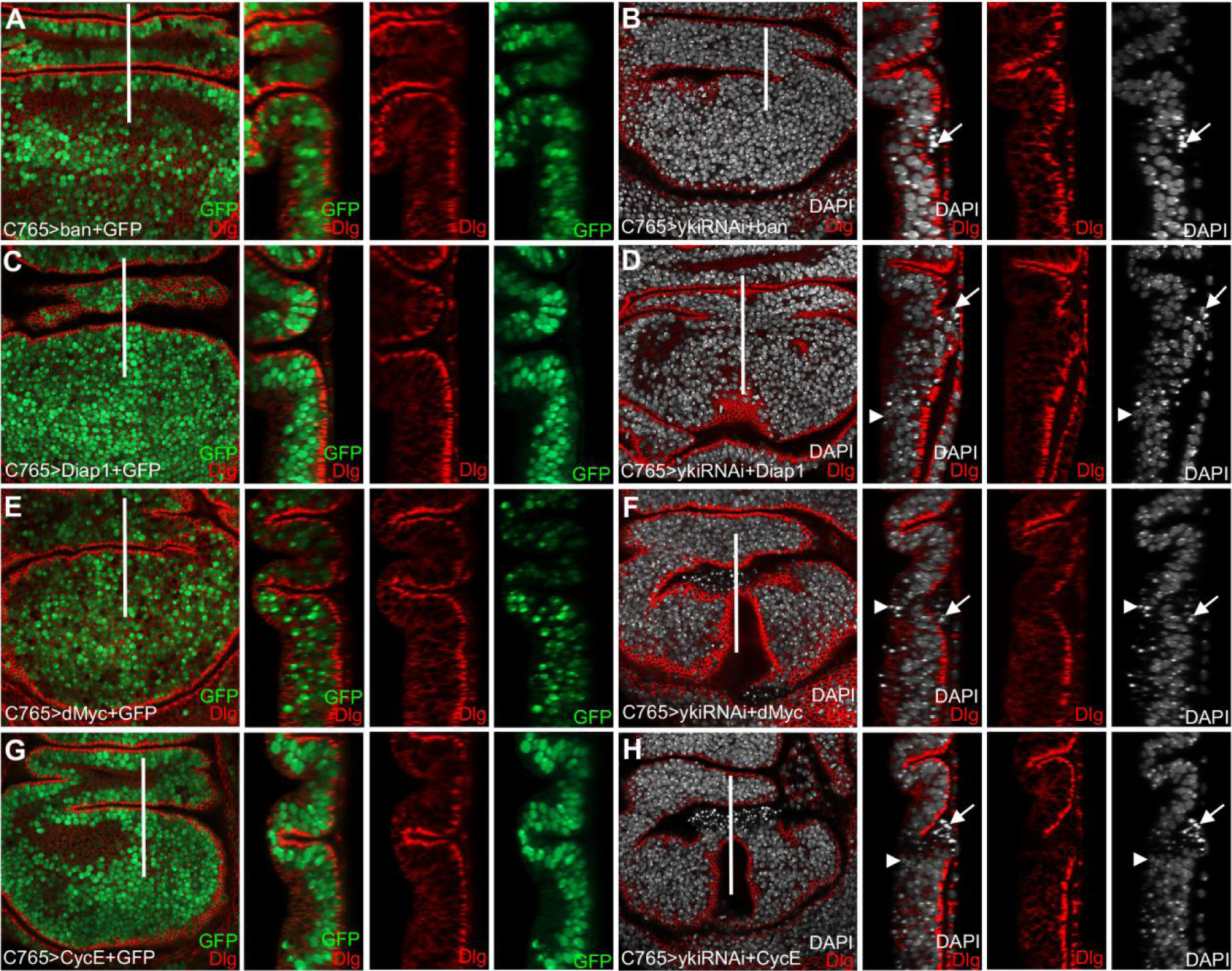
*ban* mediated BCE, but canonical Hpo-Yki targets did not mediate ACE. (A-C) ACE (arrows) induced by *ykiRNAi* was not prevented by co-expressing *ban* (B), *diap1* (D), *dMyc* (F), or *CycE* (H). (B) BCE induced by *ykiRNAi* was prevented by co-expressing *ban*. (D-H) BCE induced by *ykiRNAi* was not prevented by co-expressing *diap1* (D), *dMyc* (F), or *CycE* (H).

### Transcriptional profiles upon *yki* manipulations in the wing disc

To further investigate the mechanism of ACE induced by *ykiRNAi*, we performed RNA-Seq analysis in *ykiRNAi* expressing wing discs. In *C765*>*ykiRNAi* versus *C765*> samples, we identified 833 transcripts with more than 1.2-fold changes (false discovery rate [FDR] < 0.05), including 500 upregulated genes and 333 downregulated genes (Fig. 3A, B and Table S1). KEGG analysis of the 833 transcripts revealed the enrichment of many pathways including Hpo (Supplemental Table 2) and JNK pathways (Fig. 3C). In the *Drosophila* wing disc, ACE is associated with tumor growth and invasion (Dunn et al., 2018; Tamori et al., 2016; Vaughen and Igaki, 2016), indicating that tumor growth and invasion-involved genes are potentially related to the ACE observed. By comparing our RNA-Seq data with that produced through analyzing invasive tumors in *Drosophila* wing discs (Atkins et al., 2016), we found 90 genes that overlapped (Fig. 3E and Supplemental Table 1). Gene ontology (GO) analysis suggested that these 90 genes were enriched in carbohydrate metabolic processes, salivary gland cell autophagic cell death, cellular response to gamma radiation, epithelial cell development, and other developmental processes (Fig. 3F). Enrichment of carbohydrate metabolic processes indicates the need for macromolecule biosynthesis to support tumor growth (Külshammer et al., 2015). Salivary gland cell autophagic cell death and cellular response to gamma radiation reflect increased apoptosis. To further verify our RNA-Seq results, *yki, Ets21c* and another four genes (*kmn, puc, rpr*, and *hth*) were selected and their mRNA expression levels tested using RT-PCR (real-time, quantitative PCR). Consistent with the transcriptomic data, RT-PCR results indicate that *hth, kmn*, and *yki* were down-regulated while *puc, rpr*, and *Ets21c* were up-regulated in the *yki*RNAi wing discs (Fig. 3D and Table S2). We noticed that the expression levels of several JNK pathway components were altered in the *yki*RNAi wing discs (Fig. 3C). These results indicate that JNK signaling may be a key mediator for *yki*-dependent ACE and BCE.

**Fig. 3.**
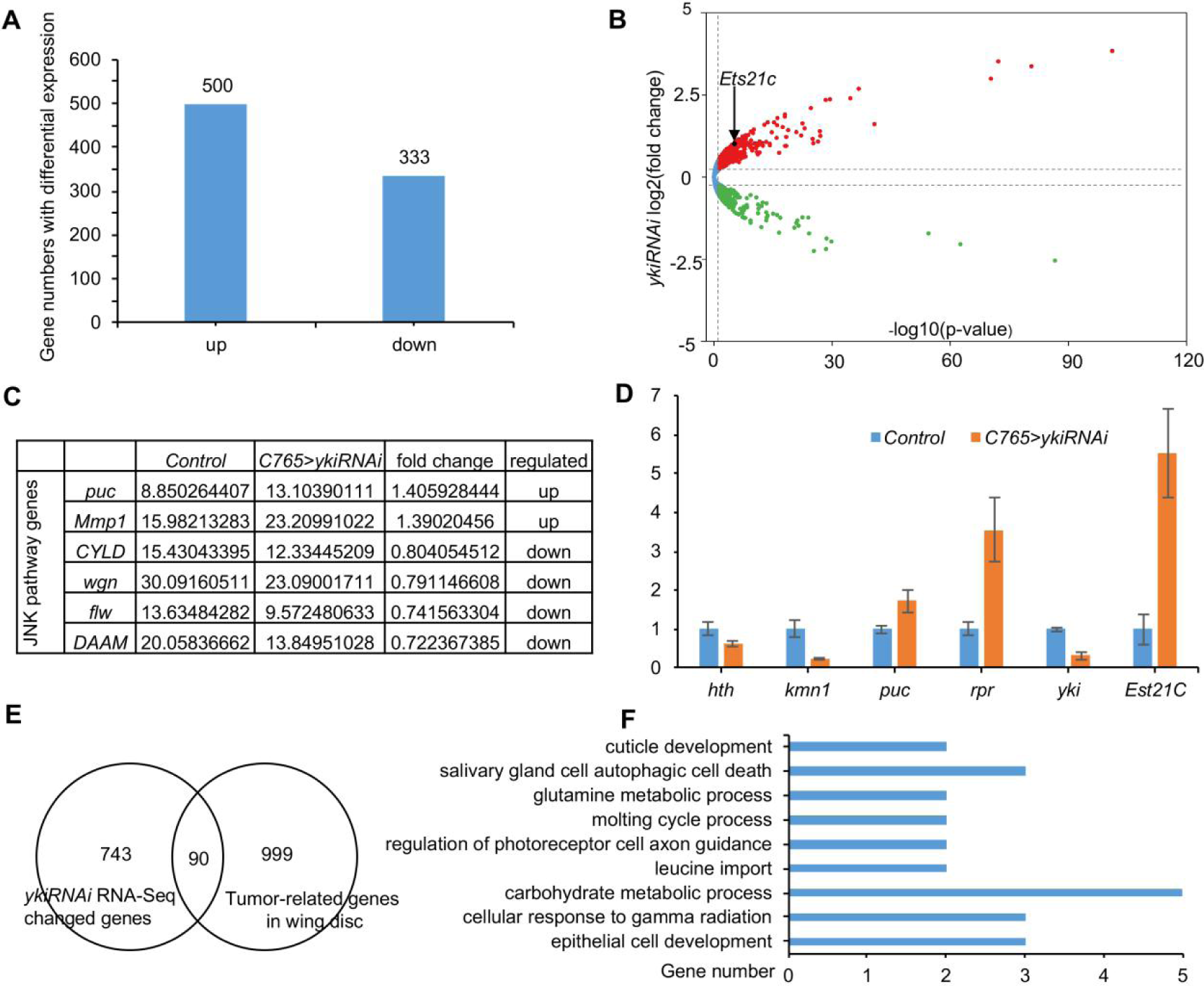
*ykiRNAi* induced gene expression profile. (A) Histogram showing the genes with altered expression in *ykiRNAi* wing discs, including 500 upregulated genes and 333 downregulated genes. (B) Distribution of genes with altered expression, black arrow pointing to the *Ets21c* gene. (C) mRNA expression alterations in six genes involved in the JNK pathway. (D) Four genes with altered expression (*hth, kmn1, puc*, and *rpr*), plus *yki* and *Ets21c*, were selected and subjected to RT-PCR. The change trends observed in the RT-PCR data were similar to that seen in the transcriptome data. (E) Venn diagram showing the 90 genes overlapping in (A) and genes changed in wing disc tumors (Atkins et al., 2016). (F) Some of these 90 genes were enriched in distinct functional GO clusters.

### Activation of JNK signaling is necessary and sufficient to induce ACE and BCE

Our transcriptome data indicated increased expression of the JNK targets *puckered* (*puc*) and *Matrix metalloproteinase 1* (*Mmp1*) (Martín-Blanco et al., 1998; Xue et al., 2007) (Fig. 3C). To confirm activation of the JNK pathway by *yki* suppression, we examined the expression levels of a *puc*-*LacZ* reporter and Mmp1 protein levels. Indeed, both *puc* and Mmp1 were upregulated in *ykiRNAi* wing discs (Fig. S5). To test whether JNK signaling is required for *ykiRNAi* induced ACE and BCE, we inhibited JNK signaling by co-expressing a dominant-negative form of JNK (*bsk*^*DN*^) (Weber et al., 2000). Both ACE and BCE were largely blocked by *bsk*^*DN*^ in *ykiRNAi* wing discs (Fig. 4A, B). The percentage of wing discs with ACE and BCE was reduced to 10%, n = 30 (Fig. S3A, C). Activation of the JNK pathway, by expressing *hep*^*CA*^ (which encodes a JNK kinase) (Glise et al., 1995), induced both ACE and BCE (Fig. 4C and Fig. S3B, D). To rule out the effect of apoptosis, we suppressed cell death by co-expressing *p35* with *hep*^*CA*^. ACE and BCE were still observed (Fig. 4D) with high frequencies (55.6 and 100%, for ACE and BCE, respectively) (Fig. S3B, D). Taken together, these results suggest that activated JNK signaling is required and sufficient to induce ACE and BCE in the *Drosophila* wing epithelia.

**Fig. 4.**
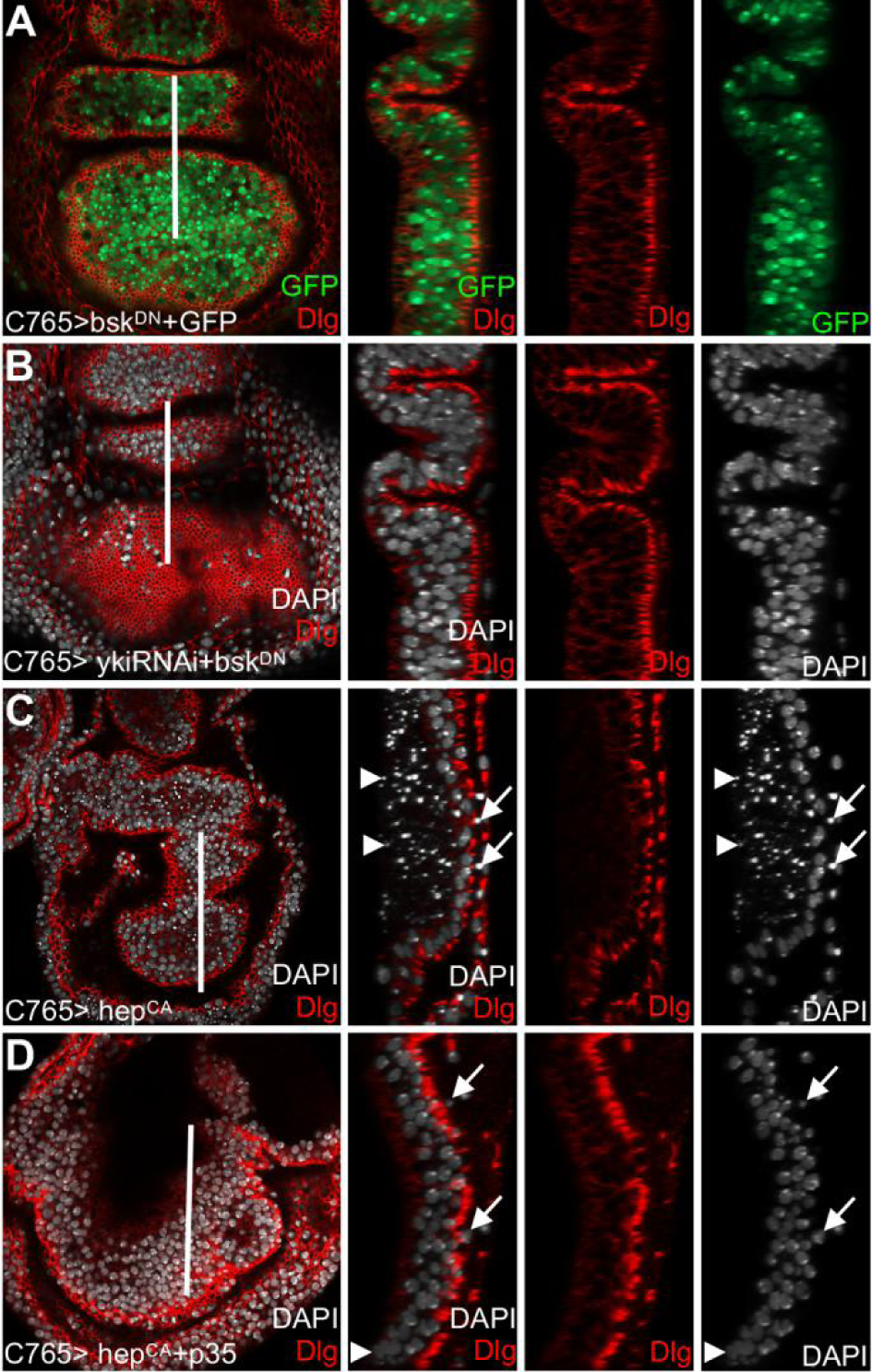
JNK signaling mediated Yki-dependent ACE and BCE. (A) Control experiment of *bsk*^*DN*^ expression. (B) Both of ACE and BCE prevented by co-expressing *bsk*^*DN*^ and *ykiRNAi*. (C) Expression of *hep*^*CA*^ induced the ACE (arrow) and BCE (arrowhead, broken cell nuclei). (D) ACE and BCE were not prevented by co-expressing *p35* and *hep*^*CA*^.

### Ets21c mediates Hpo-Yki-JNK pathway-induced ACE

To identify genes downstream of Yki-JNK that mediate ACE we performed a targeted genetic screen by either over-expressing or knocking-down the expression of candidate genes identified in the transcriptome analysis. We found that the Ets21c ETS-domain transcription factor, an ortholog of proto-oncogenes *FLI1* and *ERG*, was a key regulator of ACE. Similar to the results in *ykiRNAi* wing discs, *Ets21c* was highly upregulated in wing discs expressing *hep*^*CA*^ (Fig. S6). *UAS*-*Ets21c*^*HA*^ clones were also eliminated from the wing disc pouch (Fig. 5A), phenocopying the *ykiRNAi* clones (Fig. S1D). RT-PCR showed that *Ets21c* transcription is upregulated in *ykiRNAi* wing discs (Fig. 3D), which is consistent with our RNA-seq datasets (Fig. 3B). Ets21c expression was also examined using a GFP trap line (Jawed et al., 2017). Consistently, compared with the control (Fig. 5B, B’), *ykiRNAi* expression increased Ets21c-GFP levels (Fig. 5C, C’). These results suggest that Ets21c acts downstream of *yki*. The y-z views showed that the Ets21c-GFP level is relatively higher in cells undergoing apical extrusion (Fig. 5C, C’, yellow arrow) than that in non-extruding cells (Fig. 5C, C’). We also observed higher levels of Ets21c-GFP in apical extruding cells induced by *hep*^*CA*^ (Fig. S6B, yellow arrow). These results suggest that Ets21c might act downstream of *yki* and JNK to regulate ACE. To further confirm that Ets21c functions in Yki-JNK mediated cell extrusion, *Ets21cRNAi* was co-expressed with *ykiRNAi*. Remarkably, suppressing *Ets21c* efficiently rescued the ACE but not the BCE induced by *ykiRNAi* (Fig.5D, E, arrowhead). Co-expression of *Ets21cRNAi* with *ykiRNAi* resulted in 18.2% and 60.6% of wing discs with ACE and BCE, respectively (Fig. S3A, C, n = 33). These results suggest that *Ets21c* up-regulation mediates ACE induced by silenced *yki*. We then expressed *Ets21c*^*HA*^ to examine whether ACE can be induced. Notably, expression of *UAS*-*Ets21c*^*HA*^ was sufficient to induce ACE in 51.5% wing discs (n = 33) (Fig. 5F and Fig. S3B). Furthermore, there was no correlation between the ACE and anti-cleaved caspase-3 staining (Fig. S7), suggesting that *Ets21c*^*HA*^ induced ACE is independent of apoptosis. These results further confirm that Ets21c is a key mediator of Hpo-Yki-JNK-dependent ACE.

**Fig. 5.**
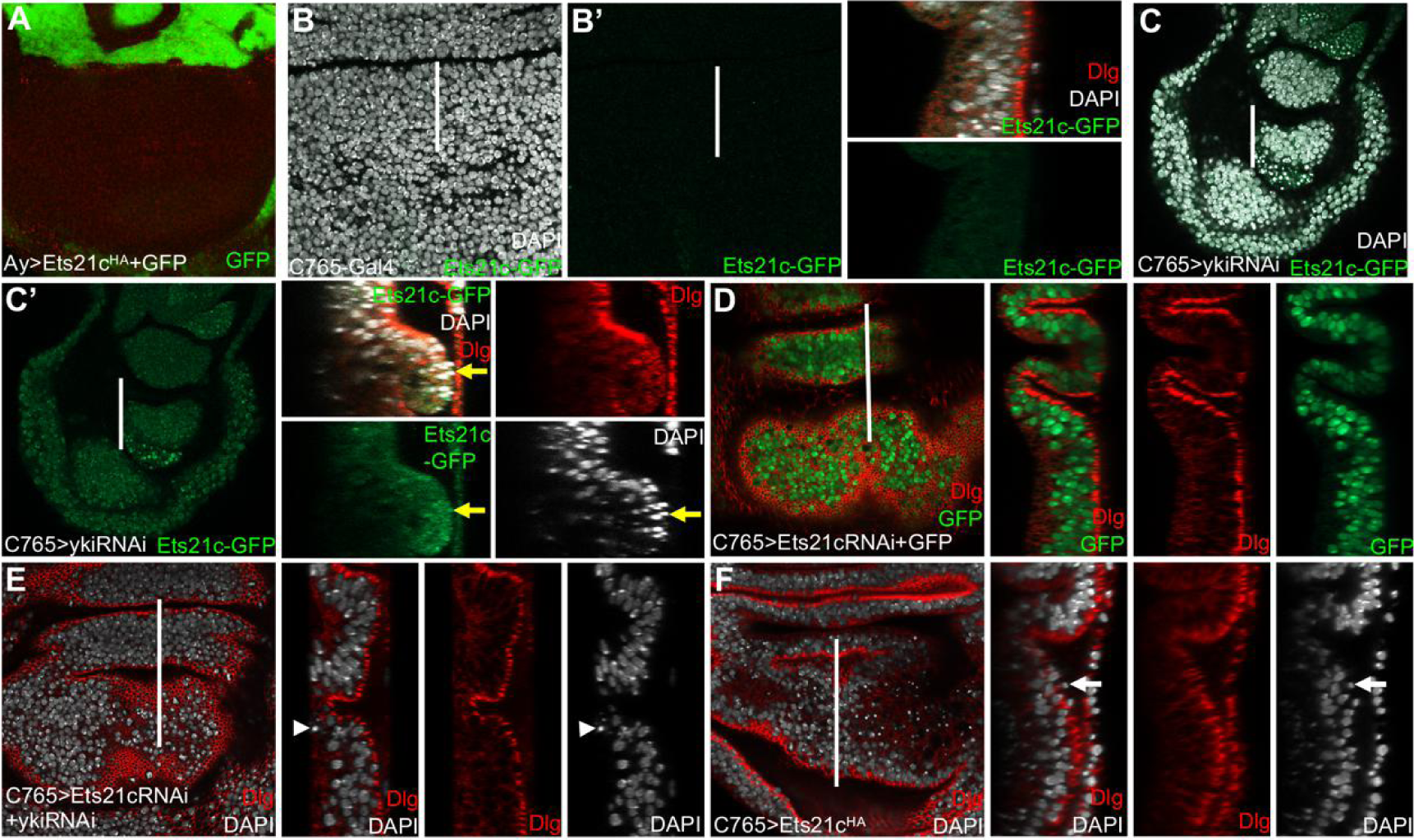
*Ets21c* mediated JNK-dependent ACE. (A) *Ets21c*^*HA*^ overexpressing clones (marked by *GFP* co-expression) were very rarely observed in the wing disc pouch. (B and B’) *Ets21c*-*GFP* level is low in control wing discs. (C and C’) *Ets21c*-*GFP* level was higher in *ykiRNAi* expressing wing discs than in the control (B). (C and C’) The *Ets21c*-*GFP* level was elevated at the position of ACE (yellow arrow). (D) Expressing *Ets21cRNAi* did not induce cell extrusion. (E) Co-expression of *Ets21cRNAi* and *ykiRNAi* largely rescued ACE, but not BCE (arrowhead). (F) Expression of *Ets21c*^*HA*^ was sufficient to induce ACE (arrow).

## DISCUSSION

Our results demonstrate that inappropriate Hpo-Yki-JNK signaling induces ACE and BCE in *Drosophila* wing disc epithelia. We also show that *ban* acts downstream of Yki to regulate BCE in the wing epithelia and that *Ets21c* acts downstream of JNK to regulate ACE in the wing epithelia.

### Activated Hpo pathway induces cell extrusion in the *Drosophila* wing disc epithelia

The Hpo pathway regulates tissue growth in *Drosophila* (Halder and Johnson, 2011). It has been reported that *yki*^*B5*^ mutant clones grow poorly in the wing and eye discs (Huang et al., 2005; Koontz et al., 2013; Qing et al., 2014). Consistent with these reports, our results showed small *ykiRNAi* and *yki*^*B5*^ mutant clones (Fig. 1B, D, and Fig. S1B, D). Cells with depleted-*yki* are extruded either apically or basally from the epithelia independently of apoptosis (Fig. 1F, Fig. S4C and Fig. 2D), indicating that cell extrusion is one explanation for the depleted-*yki* clone low recovery rate. In the *Drosophila* wing disc, overexpression of *hpo* by *MS1096*-*Gal4* and *nub*-*Gal4* dramatically decreases adult wing size (Koontz et al., 2013). Meanwhile, overexpression of *wts* by *nub*-*Gal4* also reduces the wing size (Deng et al., 2015). When we activated *hpo* and *wts* expression, using *C765*-*Gal4*, cells were intensively extruded to the lumen and the basal side of the epithelia (Fig. 1G, H). Therefore, in addition to the proliferation defect, cell extrusion is one reason for the reduced tissue size induced by Hpo pathway activation. Diap1 is decreased in the small *yki* clones (Qing et al., 2014), while co-expression of *Diap1* and *ykiRNAi* could not block ACE and BCE (Fig. 2D). These results indicate that Diap1 does not regulate cell extrusion downstream of Yki. Cells expressing low levels of dMyc were extruded basally though cell competition (Moreno and Basler, 2004). Overexpression of dMyc could not block BCE induced by silenced *yki* (Fig. 2F), indicating that other factors regulate BCE downstream of Yki. We found that *ban* could inhibit *ykiRNAi*-mediated BCE but not ACE (Fig. 2B). It is known that activated Hpo plays a role in cell-migration (Ma et al., 2017). *yki*-depleted cells migrated across the A/P boundary and extruded basally, and this cell migration was suppressed by *ban*. Our results showed that *ban* can suppress *ykiRNAi*-induced BCE in the *Drosophila* wing disc but does not regulate *ykiRNAi*-induced ACE.

In vertebrate epithelia, cells dying through apoptosis or crowding stress extrude apically into the lumen (Eisenhoffer et al., 2012; Rosenblatt et al., 2001; Slattum and Jody, 2014). The S1P–S1P2–RHO pathway regulates apoptosis-induced ACE (Gu et al., 2011; Kuipers et al., 2014). Crowding-induced ACE is dependent on regulation of the mechanosensitive ion channel PIEZIO, and subsequent activation of the S1P–S1P2 pathway (Coste et al., 2010). Several oncogenic mutations can change the normal extrusion direction from apical to basal, which leads to the high survival levels and motility of tumor cells (Slattum and Jody, 2014). In *Drosophila* epithelia, the direction of apoptotic cell extrusion is reversed with most apoptotic cells undergoing BCE (Casas-Tintó et al., 2015; Dekanty et al., 2012). Apoptosis-induced BCE is regulated by JNK signaling (Andrade and Rosenblatt, 2011). However, we know little about the mechanism of ACE in *Drosophila* epithelia, especially in normal conditions.

In *Drosophila* epithelia, apical extrusion of *scrib* mutant cells is mediated by the Slit-Robo2-Ena complex, reduced E-cadherin, and elevated Sqh levels. In normal cells, *slit, robo2*, and *ena*-overexpression only results in BCE when cell death is blocked (Vaughen and Igaki, 2016). Therefore, Slit-Robo2-Ena do not function as downstream targets of the Hpo pathway to regulate ACE. *Scrib* mutant cells activate Jak/Stat signaling and undergo ACE in the ‘tumor hotspot’ located in the dorsal hinge region of the *Drosophila* wing disc (Tamori et al., 2016). Moreover, ACE can precede M6-deficient Ras^V12^ tumor invasion following elevation of Cno/RhoA/MyoII (Dunn et al., 2018). Our RNA-Seq results revealed that the expression of Jak/Stat pathway genes and *RhoA* were not altered, indicating that ACE can be regulated by novel signaling pathways.

### JNK signaling mediates Hpo-activation-induced cell extrusion

In *Drosophila*, the JNK signaling pathway is essential for regulating cell extrusion in phenomena including wound healing, cell competition, apoptosis, and dorsal closure (Ohsawa et al., 2018). JNK signaling mediates the role of Dpp and its downstream targets in cell survival regulation in the *Drosophila* wing (Adachi-Yamada et al., 1999). Cell extrusion and retraction toward the basal side of the wing epithelia induced by the lack of Dpp activity is independent of JNK (Shen and Dahmann, 2005; Shen et al., 2010). In one case of ectopic fold formation at the A/P boundary of the *Drosophila* wing, loss of Omb activates both Yki and JNK signaling. In that case, JNK signaling induces the A/P fold by cell shortening, and Yki signaling suppresses JNK-dependent apoptosis in the folded cells (Liu et al., 2016). During cell competition induced by dMyc manipulation, JNK-dependent apoptosis mediates the death of ‘loser’ cells and their extrusion to the basal side of the epithelia. Apoptosis-induced BCE can be blocked by Diap1, which suppresses JNK-dependent apoptosis (Moreno and Basler, 2004). Taken together, these results show that JNK signaling mediates or interacts with Yki signaling in a cellular context-dependent manner during the regulation of wing epithelial morphogenesis and apoptosis.

JNK is required for the migration of *dCsk* mutant cells across the A/P boundary and their extrusion to the basal side of the epithelia (Vidal et al., 2006). *puc* encodes a JNK-specific phosphatase that provides feedback inhibition to specifically repress JNK activity. Expression of *puc* can prevent *ptc*>*dCskRNAi* cells from spreading at the A/P boundary (Vidal et al., 2006). JNK activity is also needed for *ykiRNAi* cells to invade across the wing disc A/P boundary, and co-expression of *bsk*^*DN*^ and *ykiRNAi* blocks this invasion (Ma et al., 2017). Consistent with the role JNK in BCE regulation, blocking JNK signaling by *bsk*^*DN*^ expression can prevent *ykiRNAi* cells from being extruded to the basal side of the wing epithelia (Fig. 4B). More importantly, we found that JNK activation by *hep*^*CA*^ is sufficient to induce BCE (Fig. 4C), independently of apoptosis (Fig. 4D). Furthermore, few JNK targets have been shown to regulate cell migration and BCE. An exception to this are caspases that function downstream of JNK, which can promote cell migration when activated at a mild level (Fan et al., 2020; Rudrapatna et al., 2013).

In *Drosophila* eye imaginal discs, elevated JNK signaling in *scrib* mutant cells regulates both ACE and BCE. JNK and Robo2-Ena constitute a positive feedback loop that promotes the apical and basal extrusion of *scrib* mutant cells through E-cadherin reduction. Meanwhile, in normal cells, Mmp1 upregulation when *Robo2* and *Ena* are overexpressed only induces BCE (Vaughen and Igaki, 2016). Our results showed that blocking JNK signaling could suppress ACE induced by silenced *yki* (Fig. 4B). Meanwhile, activation of JNK by *hep*^*CA*^ is sufficient to induce the extrusion of cells into the lumen (Fig. 4C). Cell debris may be trapped in the disc lumen when overexpressing *hep*^*CA*^ (Perrimon et al., 2013). We suppressed apoptosis by co-expressing *p35*, to confirm that the ACE observed was independent of cell death (Fig. 4D). Taken together, these results indicate that there are additional regulators downstream of JNK to mediate ACE in normal cells.

### *Ets21c* mediates the JNK role to regulate the ACE

E-twenty six (ETS) family transcription factors have conserved functions in metazoans (Hollenhorst et al., 2011; Sharrocks et al., 1997). These include apoptosis regulation (Sevilla et al., 1999), cell differentiation promotion (Taylor et al., 1997), cell fate regulation (Miley et al., 2004), and cellular senescence (Ohtani et al., 2001). *Ets21c* is a member of the ETS-domain transcription factor family and it is the ortholog of the human proto-oncogenes *FLI1* and *ERG* (Mundorf et al., 2019). In *Drosophila* eye imaginal discs, 30-fold increased *Ets21c* expression is induced by *Ras*^*V12*^ and *Eiger*, an activator of JNK (Toggweiler et al., 2016). In the *Drosophila* adult midgut, *Ets21c* expression is increased when JNK is activated by the JNK kinase *hep* (Mundorf et al., 2019). Ets21c also can promote tumor growth downstream of the JNK pathway (Külshammer et al., 2015; Toggweiler et al., 2016). Our results have confirmed that Ets21c functions downstream of JNK. Indeed, we have shown that the Ets21c-GFP level is elevated following JNK activation (Fig. S6A). Expression of *Ets21c*^*HA*^ is sufficient to induce ACE (Fig. 5F) and silencing of *Ets21c* is sufficient to rescue *ykiRNAi*-induced ACE in the wing discs (Fig. 5E). However, the mechanism through which *yki* regulates JNK-Ets21c remains to be determined.

ACE promotes polarity-impaired cells to grow into tumors (Tamori et al., 2016; Vaughen and Igaki, 2016). Therefore, it is possible that Ets21c can promote Hpo-Yki-JNK related tumorigenesis by facilitating ACE. Yki/YAP gain-of-function promotes cancer cell invasion in non-small-cell lung cancer (Yang et al., 2010), neoplastic transformation (Yang et al., 2013), uveal melanoma (Yu et al., 2014), and pancreatic cancer (Yang et al., 2015). Additionally, Yki/YAP loss-of-function helps tumor cells to escape from apoptosis in hematologic malignancies, including multiple myeloma, lymphoma, and leukemia (Cottini et al., 2014). Consistent with the latter role, Yki suppresses cell extrusion from the *Drosophila* wing epithelia by suppressing Ets21c (Fig. 5E). Therefore, the role of Ets21c in Hpo-Yki related tumor models should be further examined.

## MATERIALS AND METHODS

### *Drosophila* genetics

The following transgenes were used in this study: *C765*-*Gal4* (Nellen et al., 1996), *UAS*-*bsk*^*DN*^ (Weber et al., 2000), *UAS*-*ban* (Ying and Padgett, 2012), *puc*-*lacZ* (Martín-Blanco et al., 1998), *FRT42D*-*yki*^*B5*^ and *FRT42D* (gifts from Z.H. Li), *UAS*-*Diap1* (gift from A. Bergmann), *UAS*-*hippo* and *UAS*-*wts* (gifts from L. Xue), *UAS*-*GFP* (BL#4775), *UAS*-*CycE* (BL#30725), *UAS*-*dMyc* (BL#9674), *UAS*-*p35* (BL#5073), *UAS*-*hep*^*CA*^ (BL#38639), *Ets21c*-*GFP* (BL#38639), *UAS*-*Ets21cRNAi* (VDRC#106153), *UAS*-*Ets21c*^*HA*^ (FlyORF#F000624), *UAS*-*ykiRNAi* (THU#0579), and *UAS*-*ykiRNAi* (THU#3074).

### Transgene expression and Clonal induction

Larvae were raised at 25°C. For efficient expression of *RNAi* and *UAS* transgenes driven by the *C765*-*Gal4*, larvae were raised at 29°C. To generate *UAS*-*ykiRNAi* and *UAS*-*Ets21c* clones, larvae of genotype *y w hs*-*Flp*/+; *act5c*>*CD2*>*Gal4, UAS*-*GFP*/+; *UAS*-*ykiRNAi*/+ and *y w hs*-*Flp*/+; *act5c*>*CD2*>*Gal4, UAS*-*GFP*/+; *UAS*-*Ets21c*/+ were subjected to heat shock at 35°C for 30 minutes. To generate *yki*^*B5*^ MARCM clones, larvae of genotype *y w hs*-*flp, tub*-*Gal4, UAS*-*GFP*/+; *FRT42D, Tub*-*Gal80*/*FRT42D yki*^*B5*^ were subjected to heat shock for 1 hour at 37.5°C. Larval genotypes of every figures were listed in Table S5.

### Immunohistochemistry

*Drosophila* wing imaginal discs were dissected from middle to late third instar larvae and antibody staining was performed according to standard procedures. The primary antibodies used were: mouse anti-Dlg (1:10, Developmental Studies Hybridoma Bank, DSHB); rabbit anti-cleaved caspase-3 (1:200, Cell Signaling); mouse anti-Mmp1 (1:10, DSHB); and mouse anti-β-galactosidase, (1:2000, Promega). Secondary antibodies (diluted 1:200) used were anti-mouse Cy3, anti-rabbit Cy3, and anti-rat Cy3 (Jackson Immuno Research). The cell membrane was stained using rhodamine-phalloidin (1:1000, Sigma). Cell nuclei were stained with DAPI (1:200000, Sigma). Images were collected using Leica SP8 and Zeiss LSM 800 confocal microscopes. The figures were assembled in Adobe Photoshop CC with minor image adjustments (brightness and/or contrast).

### RNA-Seq

*C765*>*ykiRNAi* wing discs were dissected as test samples, and *C765*> wing discs were dissected as controls. For each sample, 100 wing discs were dissected in PBS on ice. Total RNA was prepared with Trizol regent (Invitrogen™, Code No. 12183555) following the recommendations of the manufacturer. RNA quality was verified by agarose gel electrophoresis and by Nanodrop 2,000. RNA-Seq libraries were sequenced on an Illumina Hiseq platform. Raw reads were processed to clean data using in-house perl scripts. Clean data were mapped to the *Drosophila* genome using TopHat v2.0.12 (Kim et al., 2013). Analysis of differential expression in test and control samples was performed using the DESeq2 R package. Genes with DESeq2 adjusted P-values < 0.01 were considered to be differentially expressed. KOBAS software was used to test the statistical enrichment of differentially expressed genes in KEGG pathways (http://www.genome.jp/kegg/). We also used DAVID to test the GO functional enrichment of differentially expressed genes (https://david.ncifcrf.gov).

### Quantitative real-time PCR

cDNA was reversed transcribed from RNA using the Takara PrimeScript™ RT reagent Kit with gDNA Eraser (Code No. RR047A). qRT-PCR was performed using the PerfectStart™ Green qPCR SuperMix (+Dye II) in the QuantStudio 6 Flex platform and each reaction was repeated in triplicate. Transcription levels were normalized to those of *act*. Data were calculated using Microsoft Excel.

The gene-specific primer sequences used are showed in Supplemental Table 4.

## Acknowledgements

We thank Dr. Xue L., Dr. Z.H. Li, Dr. A. Bergmann Bloomington Stock Center, Vienna Drosophila RNAi Center and Tsinghua University Stock Center for fly stocks, and Dr. N. Jiang and Dr. L. Qi for the confocal facilities.

## Author contributions

XA performed the experiments. XA, JS, JZ, and DW designed the experiments, analyzed and interpreted the data, and wrote the manuscript.

## Compliance with ethical standards

### Supplementary figures

Supplementary Figures and Tables are listed in detail in the Supplementary Materials.

## References

Adachi-Yamada, T., Fujimura-Kamada, K., Nishida, Y. and Matsumoto, K. (1999). Distortion of proximodistal information causes JNK-dependent apoptosis Drosophila wing. Nature 400, 166–169.

Andrade, D. and Rosenblatt, J. (2011). Apoptotic regulation of epithelial cellular extrusion. Apoptosis 16, 491–501.

Atkins, M., Potier, D., Romanelli, L., Jacobs, J., Mach, J., Hamaratoglu, F., Aerts, S. and Halder, G. (2016). An Ectopic Network of Transcription Factors Regulated by Hippo Signaling Drives Growth and Invasion of a Malignant Tumor Model. Curr. Biol. 26, 2101–2113.

Casas-Tintó, S., Lolo, F.-N. and Moreno, E. (2015). Active JNK-dependent secretion of Drosophila Tyrosyl-tRNA synthetase by loser cells recruits haemocytes during cell competition. Nat. Commun. 6, 10022.

Coste, B., Mathur, J., Schmidt, M., Earley, T. J., Ranade, S., Petrus, M. J., Dubin, A. E. and Patapoutian, A. (2010). Piezo1 and Piezo2 Are Essential Components of Distinct Mechanically Activated Cation Channels. Science 330, 55–60.

Cottini, F., Hideshima, T., Xu, C., Sattler, M., Dori, M., Agnelli, L., Hacken, E. t., Bertilaccio, M. T., Antonini, E., Neri, A., et al. (2014). Rescue of Hippo coactivator YAP1 triggers DNA damage-induced apoptosis in hematological cancers. Nat. Med. 20, 599–606.

Dekanty, A., Barrio, L., Muzzopappa, M., Auer, H. and Milán, M. (2012). Aneuploidy-induced delaminating cells drive tumorigenesis in Drosophila epithelia. Proc. Natl. Acad. Sci. USA 109, 20549–20554.

Deng, H., Wang, W., Yu, J., Zheng, Y., Yun, Q. and Pan, D. (2015). Spectrin regulates Hippo signaling by modulating cortical actomyosin activity. elife 4, e06567.

Dong, J., Feldmann, G., Huang, J., Wu, S., Zhang, N., Comerford, S. A., Gayyed, Mariana F., Anders, R. A., Maitra, A. and Pan, D. (2007). Elucidation of a Universal Size-Control Mechanism in Drosophila and Mammals. Cell 130, 1120–1133.

Dunn, B. S., Rush, L., Lu, J. and Xu, T. (2018). Mutations in the Drosophila tricellular junction protein M6 synergize with RasV12 to induce apical cell delamination and invasion. Proc. Natl. Acad. Sci. USA 115, 8358–8363.

Eisenhoffer, G. T., Loftus, P. D., Yoshigi, M., Otsuna, H., Chien, C.-B., Morcos, P. A. and Rosenblatt, J. (2012). Crowding induces live cell extrusion to maintain homeostatic cell numbers in epithelia. Nature 484, 546–549.

Fan, W., Luo, D., Zhang, J., Wang, D. and Shen, J. (2020). Vestigial suppresses apoptosis and cell migration in a manner dependent on the level of JNK/Caspase signaling in the Drosophila wing disc. Insect Sci., DOI: 10.1111/1744-7917.12762.

Feng, X., Degese, M. S., Iglesias-Bartolome, R., Vaque, J. P., Molinolo, A. A., Rodrigues, M., Zaidi, M. R., Ksander, B. R., Merlino, G., Sodhi, A., et al. (2014). Hippo-independent activation of YAP by the GNAQ uveal melanoma oncogene through a trio-regulated rho GTPase signaling circuitry. Cancer Cell 25, 831–845.

Glise, B., Bourbon, H. G. and Noselli, S. (1995). hemipterous encodes a novel drosophila MAP kinase kinase, required for epithelial cell sheet movement. Cell 83, 451–461.

Gu, Y., Forostyan, T., Sabbadini, R. A. and Rosenblatt, J. (2011). Epithelial cell extrusion requires the sphingosine-1-phosphate receptor 2 pathway. J. Cell Biol. 193, 667–676.

Gu, Y. and Rosenblatt, J. (2012). New emerging roles for epithelial cell extrusion. Curr. Opin. Cell Biol. 24, 865–870.

Gudipaty, S. A. and Rosenblatt, J. (2016). Epithelial cell extrusion: Pathways and pathologies. Semin. Cell Dev. Biol. 67, 132–140.

Halder, G. and Johnson, R. L. (2011). Hippo signaling: growth control and beyond. Development 138, 9–22.

Harrison, D. A. and Perrimon, N. (1993). Simple and efficient generation of marked clones in Drosophila. Curr. Biol. 3, 424–433.

Hay, B. A., Wassarman, D. A. and Rubin, G. M. (1995). Drosophila homologs of baculovirus inhibitor of apoptosis proteins function to block cell death. Cell 83, 1253–1262.

Herrera, S. C., Martín, R. and Morata, G. (2013). Tissue Homeostasis in the Wing Disc of Drosophila melanogaster: Immediate Response to Massive Damage during Development. PLoS Genet. 9, e1003446.

Hollenhorst, P. C., McIntosh, L. P. and Graves, B. J. (2011). Genomic and Biochemical Insights into the Specificity of ETS Transcription Factors. Annu. Rev. Biochem. 80, 437–471.

Huang, J., Wu, S., Barrera, J., Matthews, K. and Pan, D. (2005). The Hippo signaling pathway coordinately regulates cell proliferation and apoptosis by inactivating Yorkie, the Drosophila Homolog of YAP. Cell 122, 421–434.

Khan, S. J., Abidi, S. N. F., Skinner, A., Tian, Y. and Smith-Bolton, R. K. (2017). The Drosophila Duox maturation factor is a key component of a positive feedback loop that sustains regeneration signaling. PLoS Genet. 13, e1006937.

Kim, D., Pertea, G., Trapnell, C., Pimentel, H., Kelley, R. and Salzberg, S. L. (2013). TopHat2: accurate alignment of transcriptomes in the presence of insertions, deletions and gene fusions. Genome Biol. 14, R36.

Knust, E. and Bossinger, O. (2002). Composition and formation of intercellular junctions in epithelial cells. Science 298, 1955–1959.

Koontz, L. M., Liu-Chittenden, Y., Yin, F., Zheng, Y., Yu, J., Huang, B., Chen, Q., Wu, S. and Pan, D. (2013). The Hippo effector Yorkie controls normal tissue growth by antagonizing scalloped-mediated default repression. Dev. Cell 25, 388–401.

Kuipers, D., Mehonic, A., Kajita, M., Peter, L., Fujita, Y., Duke, T., Charras, G. and Gale, J. E. (2014). Epithelial repair is a two-stage process driven first by dying cells and then by their neighbours. J. Cell Sci. 127, 1229–1241.

Külshammer, E., Mundorf, J., Kilinc, M., Frommolt, P., Wagle, P. and Uhlirova, M. (2015). Interplay among Drosophila transcription factors Ets21c, Fos and Ftz-F1 drives JNK-mediated tumor malignancy. Dis. Models Mech. 8, 1279–1293.

Lampaya, M. E. and Basler, K. (2004). dMyc Transforms Cells into Super-Competitors. Cell 117, 117–129.

Lee, T. and Luo, L. (2001). Mosaic analysis with a repressible cell marker (MARCM) for Drosophila neural development. Trends Neurosci. 24, 251–254.

Liu, S., Sun, J., Wang, D., Pflugfelder, G. O. and Shen, J. (2016). Fold formation at the compartment boundary of Drosophila wing requires Yki signaling to suppress JNK dependent apoptosis. Sci. Rep. 6, 38003.

Ma, X., Wang, H., Ji, J., Xu, W., Sun, Y., Li, W., Zhang, X., Chen, J. and Xue, L. (2017). Hippo signaling promotes JNK-dependent cell migration. Proc. Natl. Acad. Sci. USA 114, 1934–1939.

Marinari, E., Mehonic, A., Curran, S., Gale, J., Duke, T. and Baum, B. (2012). Live-cell delamination counterbalances epithelial growth to limit tissue overcrowding. Nature 484, 542–545.

Martín-Blanco, E., Gampel, A., Ring, J., Virdee, K., Kirov, N., Tolkovsky, A. M. and Martinez-Arias, A. (1998). puckered encodes a phosphatase that mediates a feedback loop regulating JNK activity during dorsal closure in Drosophila. Genes Dev. 12, 557–570.

Miley, G. R., Fantz, D. A., Glossip, D., Lu, X., Saito, R. M., Palmer, R. E., Inoue, T., van den Heuvel, S., Sternberg, P. W. and Kornfeld, K. (2004). Identification of Residues of the Caenorhabditis elegans LIN-1 ETS Domain That Are Necessary for DNA Binding and Regulation of Vulval Cell Fates. Genetics 167, 1697–1709.

Mundorf, J., Donohoe, C. D., McClure, C. D., Southall, T. D. and Uhlirova, M. (2019). Ets21c Governs Tissue Renewal, Stress Tolerance, and Aging in the Drosophila Intestine. Cell Rep. 27, 3019–3033.

Nellen, D., Burke, R., Struhl, G. and Basler, K. (1996). Direct and Long-Range Action of a DPP Morphogen Gradient. Cell 85, 357–368.

Neto-Silva, R. M., Beco, S. D. and Johnston, L. A. (2010). Evidence for a Growth-Stabilizing Regulatory Feedback Mechanism between Myc and Yorkie, the Drosophila Homolog of Yap. Dev. Cell 19, 507–520.

Nolo, R., Morrison, C. M., Tao, C., Zhang, X. and Halder, G. (2006). The bantam MicroRNA Is a Target of the Hippo Tumor-Suppressor Pathway. Curr. Biol. 16, 1895–1904.

Ohsawa, S., Vaughen, J. and Igaki, T. (2018). Cell Extrusion: A Stress-Responsive Force for Good or Evil in Epithelial Homeostasis. Dev. Cell 44, 284–296.

Ohtani, N., Zebedee, Z., Huot, T. J. G., Stinson, J. A., Sugimoto, M., Ohashi, Y., Sharrocks, A. D., Peters, G. and Hara, E. (2001). Opposing effects of Ets and Id proteins on p16INK4a expression during cellular senescence. Nature 409, 1067–1070.

Pagliarini, R. A. and Xu, T. (2003). A Genetic Screen in Drosophila for Metastatic Behavior. Science 302, 1227–1231.

Pan, D. (2010). The Hippo Signaling Pathway in Development and Cancer. Dev. Cell 19, 491–505.

Praskova, M., Xia, F. and Avruch, J. (2008). MOBKL1A/MOBKL1B phosphorylation by MST1 and MST2 inhibits cell proliferation. Curr. Biol. 18, 311–321.

Qing, Y., Yin, F., Wang, W., Zheng, Y., Guo, P., Schozer, F., Deng, H. and Pan, D. (2014). The Hippo effector Yorkie activates transcription by interacting with a histone methyltransferase complex through Ncoa6. Elife 3, e02564.

Rosenblatt, J., Raff, M. C. and Cramer, L. P. (2001). An epithelial cell destined for apoptosis signals its neighbors to extrude it by an actin- and myosin-dependent mechanism. Curr. Biol. 11, 1847–1857.

Rudrapatna, V. A., Bangi, E. and Cagan, R. L. (2013). Caspase signalling in the absence of apoptosis drives Jnk-dependent invasion. EMBO Rep. 14, 172–177.

Schroeder, M. C. and Halder, G. (2012). Regulation of the Hippo pathway by cell architecture and mechanical signals. Semin. Cell Dev. Biol. 23, 803–811.

Sevilla, L., Aperlo, C., Dulic, V., Chambard, J. C., Boutonnet, C., Pasquier, O., Pognonec, P. and Boulukos, K. E. (1999). The Ets2 Transcription Factor Inhibits Apoptosis Induced by Colony-Stimulating Factor 1 Deprivation of Macrophages through a Bcl-xL-Dependent Mechanism. Mol. Cell. Biol. 19, 2624–2634.

Sharrocks, A. D., Brown, A. L., Ling, Y. and Yates, P. R. (1998). The ETS-domain transcription factor family. Int. J. Biochem. Cell Biol. 29, 1371–1387.

Shen, J. and Dahmann, C. (2005). Extrusion of cells with inappropriate Dpp signaling from Drosophila wing disc epithelia. Science 307, 1789–1790.

Shen, J., Dahmann, C. and Pflugfelder, G. O. (2010). Spatial discontinuity of Optomotor-blind expression in the Drosophila wing imaginal disc disrupts epithelial architecture and promotes cell sorting. BMC Dev. Biol. 10, 23.

Shen, J., Lu, J., Sui, L., Wang, D., Yin, M., Hoffmann, I., Legler, A. and Pflugfelder, G. O. (2014). The orthologous Tbx transcription factors Omb and TBX2 induce epithelial cell migration and extrusion in vivo without involvement of matrix metalloproteinases. Oncotarget 5, 11998–12015.

Slattum, G. M. and Jody, R. (2014). Tumour cell invasion: an emerging role for basal epithelial cell extrusion. Nat. Rev. Cancer 14, 495–501.

Tamori, Y., Suzuki, E. and Deng, W. (2016). Epithelial Tumors Originate in Tumor Hotspots, a Tissue-Intrinsic Microenvironment. PLoS Biol. 14, e1002537.

Tapon, N., Harvey, K. F., Bell, D. W., Wahrer, D. C. R., Schiripo, T. A., Haber, D. A. and Hariharan, I. K. (2002). salvador Promotes Both Cell Cycle Exit and Apoptosis in Drosophila and Is Mutated in Human Cancer Cell Lines. Cell 110, 467–478.

Taylor, J. M., Dupont-Versteegden, E. E., Davies, J. D., Hassell, J. A., Houlé, J. D., Gurley, C. M. and Peterson, C. A. (1997). A role for the ETS domain transcription factor PEA3 in myogenic differentiation. Mol. Cell. Biol. 17, 5550–5558.

Thompson, B. J. and Cohen, S. M. (2006). The Hippo pathway regulates the bantam microRNA to control cell proliferation and apoptosis in Drosophila. Cell 126, 767–774.

Toggweiler, J., Willecke, M. and Basler, K. (2016). The transcription factor Ets21C drives tumor growth by cooperating with AP-1. Sci. Rep. 6, 34725.

Udan, R. S., Kango-Singh, M., Nolo, R., Tao, C. and Halder, G. (2003). Hippo promotes proliferation arrest and apoptosis in the Salvador/Warts pathway. Nat. Cell Biol. 5, 914–920.

Vaughen, J. and Igaki, T. (2016). Slit-Robo Repulsive Signaling Extrudes Tumorigenic Cells from Epithelia. Dev. Cell 39, 683–695.

Vidal, M., Larson, D. E. and Cagan, R. (2006). Csk-Deficient Boundary Cells Are Eliminated from Normal Drosophila Epithelia by Exclusion, Migration, and Apoptosis. Dev. Cell 10, 33–44.

Wang, Y., Dong, Q., Zhang, Q., Li, Z., Wang, E. and Qiu, X. (2010). Overexpression of yes-associated protein contributes to progression and poor prognosis of non-small-cell lung cancer. Cancer Sci. 101, 1279–1285.

Weber, U., Paricio, N. and Mlodzik, M. (2000). Jun mediates Frizzled-induced R3/R4 cell fate distinction and planar polarity determination in the Drosophila eye. Development 127, 3619–3629.

Wu, S., Huang, J., Dong, J. and Pan, D. (2003). hippo Encodes a Ste-20 Family Protein Kinase that Restricts Cell Proliferation and Promotes Apoptosis in Conjunction with salvador and warts. Cell 114, 445–456.

Xue, L., Igaki, T., Kuranaga, E., Kanda, H., Miura, M. and Xu, T. (2007). Tumor Suppressor CYLD Regulates JNK-Induced Cell Death in Drosophila. Dev. Cell 13, 446–454.

Yang, S., Zhang, L., Liu, M., Chong, R., Ding, S., Chen, Y. and Dong, J. (2013). CDK1 Phosphorylation of YAP Promotes Mitotic Defects and Cell Motility and Is Essential for Neoplastic Transformation. Cancer Res. 73, 6722–6733.

Yang, S., Zhang, L., Purohit, V., Shukla, S. K., Chen, X., Yu, F., Fu, K., Chen, Y., Solheim, J., Singh, P. K., et al. (2015). Active YAP promotes pancreatic cancer cell motility, invasion and tumorigenesis in a mitotic phosphorylation-dependent manner through LPAR3. Oncotarget 6, 36019–36031.

Ying, L. and Padgett, R. W. (2012). bantam Is Required for Optic Lobe Development and Glial Cell Proliferation. PLoS One 7, e32910.

Yu, F., Luo, J., Mo, J., Liu, G., Kim, Y. C., Meng, Z., Zhao, L., Peyman, G., Ouyang, H., Jiang, W., et al. (2014). Mutant Gq/11 promote uveal melanoma tumorigenesis by activating YAP. Cancer Cell 25, 822–830.

Zhang, L., Ren, F., Zhang, Q., Chen, Y., Wang, B. and Jiang, J. (2008). The TEAD/TEF Family of Transcription Factor Scalloped Mediates Hippo Signaling in Organ Size Control. Dev. Cell 14, 377–387.

